# BioSANS: A Software Package for Symbolic and Numeric Biological Simulation

**DOI:** 10.1101/2021.08.17.456661

**Authors:** Erickson Fajiculay, Chao-Ping Hsu

**Affiliations:** Institute of Chemistry, Academia Sinica, Taipei, 11529, Taiwan; Bioinformatics Program, Institute of Information Science, Taiwan International Graduate Program, Academia Sinica, Taipei, 11529, Taiwan; Institute of Bioinformatics and Structure Biology, National Tsinghua University, 30044, Taiwan; Physics Division, National Center for Theoretical Sciences, Taipei, 10617, Taiwan; Genome and Systems Biology Degree program, National Taiwan University, Taipei, 10617, Taiwan

## Abstract

Modeling biochemical systems can provide insights into behaviors that are difficult to observe or understand. It requires software, programming, and understanding of the system to build a model and study it. Softwares exist for systems biology modeling, but most support only certain types of modeling tasks. Desirable features including ease in preparing input, symbolic or analytical computation, parameter estimation, graphical user interface, and systems biology markup language (SBML) support are not seen concurrently in one software package. In this study, we developed a python-based software that supports these features, with both deterministic and stochastic propagations. The software can be used by graphical user interface, command line, or as a python import. We also developed a semi-programmable and intuitively easy topology input method for the biochemical reactions. We tested the software with semantic and stochastic SBML test cases. Tests on symbolic solution and parameter estimation were also included. The software we developed is reliable, well performing, convenient to use, and compliant with most of the SBML tests. So far it is the only systems biology software that supports symbolic, deterministic, and stochastic modeling in one package that also features parameter estimation and SBML support. This work offers a comprehensive set of tools and allows for better availability and accessibility for studying kinetics and dynamics in biochemical systems.

## 1.0 Introduction

The complex nature of biological systems often hinders a full understanding of behavior, an area in which mathematical models and simulation can help [1]. Clues from experiments are limited to the details of the sub-systems considered and can be difficult to interpret. Computer simulation allows for formulating a working model that can help explain experimental observations [2,3]. It can provide links between observations and unknowns in terms of a mathematical expression or numerical values, offering qualitative or quantitative insights. Good models can give testable predictions that can be used to evaluate the applicability, scope, and limitation of the model [4,5]. No model can fully account for every detail of a system, but some can provide important aspects of the system within its scope [6].

Interest in modeling biological processes has been increasing [7]. Models of processes that constitute a network of interacting molecules are of particular interest [8–12]. Algorithms and theories dedicated to modeling gene expressions have been developed [13–15]. Modeling and experimental studies related to gene expression, biological clock, and diseases are now common [16–20].

Simulating a model is not difficult, but for a complicated model or for elaborated analyses and tests with models, scripting or programming skills are necessary. To simulate a model, high-level programming tools such as Matlab or Python are often used. Complicated models may require advance programming skills and deep understanding of the domain concepts involved. Software exists for modeling biological systems; most require basic scripting [21–28]. Some do not require coding for basic simulations but have limited power in the graphical user interface (GUI) [29]. For most software, the propensity expression needs to be manually typed by the user. This is sometimes time-consuming and a hindrance to usability. Most systems-biology software packages support only certain types of modeling tasks. As far as we know, no systems biology software provides symbolic computation capability without the need to declare variables and write ordinary differential equation (ODE) expressions. Most are not user friendly, do not allow for easy-to-prepare input, do not support parameter estimation and do not provide a GUI. Some do not provide systems biology markup language (SBML) support and others have limited or minimal SBML supported features. Therefore, a software package addressing the above limitations is highly desirable because it allows access to mathematical modeling by a much broader range of users.

In this study, we took advantage of Python libraries for both numeric and symbolic computation and developed a program that can make systems biology analysis available to non-domain experts. In doing so, we provide a platform that can support a wide variety of modeling tasks. Currently, the software supports model construction, symbolic computation, propagation, analysis, and parameter estimation and also supports SBML. A semi-programmable topological model input was developed, for easy construction and understanding. We also provide a GUI. The algorithms are available in Python import format for expert users and also via a command line using our novel structured simulation language (SSL), which is very similar to MySQL commands in terms of readability. We tested the software in simulating systems of various sizes, and it was found reliable, practical, and easy to use. We provide an invaluable research tool for model simulations by domain experts from systems biology or chemistry.

## 2.0 Modeling scheme supported in BioSANS

The process of a typical modeling task generally involves model construction, propagation and analysis. Fig 1 lists these processes with functionalities provided in the developed software, BioSANS. The first step is to build a topological model and to establish the set of differential equations based on the topology of the model. For the initial model, temporary parameter values are required. These values can be based on databases, experimental design, physical intuition, and known ground truth. If sufficient data are available, parameter estimation may help tune the parameters. If the ODE is analytically solvable, species analytical expression as a function of time, initial conditions, and rate constants can be computed. If not, numerical integration techniques can be used to propagate the trajectories. Propagation can be deterministic or stochastic and compared to experimental data to gain some physical insights.

**Fig 1.**
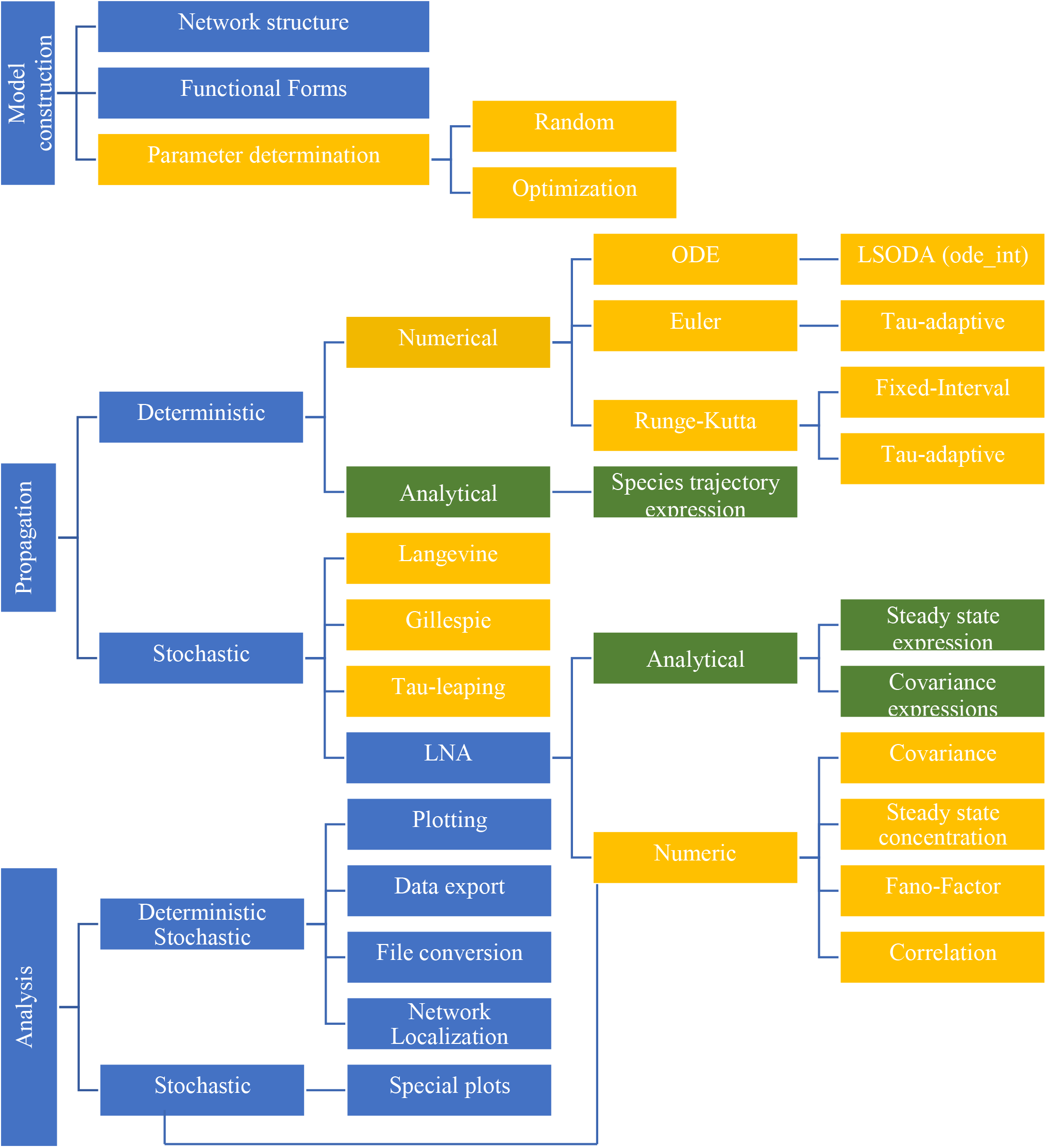
Schematic diagram showing some of the modeling tasks available in BioSANS. The basic modeling scheme includes model construction, propagation, and analysis. Except for blue colors, the same color in different branches indicates similar types of analysis under a common category. Green shaded boxes are for analytical expressions, and orange shaded boxes are for numerical results.

Linear noise approximation (LNA) is a convenient way to study noise in the system [30– 32]. Derivation of analytical expression utilizing the LNA is supported in BioSANS and can be compared to the stochastic results. This allows for gaining physical insights into the noise effects in the system as a function of physical variables.

Once the simulation is performed and the analytical expressions are calculated, further analyses such as plotting, calculation of statistics, etc. can be performed to judge the result and to check consistencies with known ground truth. If the model has inconsistencies and/or fails to account for experimental observations, some of the hypotheses put forth may be proved wrong, and the model may need some modification. A cycle from gathering experimental data, model construction and simulation is needed to ensure consistency between the model and observations. A full list of modeling tasks and workflow to be followed in using BioSANS on a case-to-case basis is included in supporting information.

## 3.0 Description of algorithms and implementation

The codes in BioSANS are written in Python [33], which only needs a user-defined topology file that is intuitive to construct based on basic chemistry knowledge. The software algorithm grabs data from the topology file and calculates the stoichiometric matrix and propensity vector. It then sets up the ordinary differential equation for most typical types of simulations. This process is handled by building a dictionary of species, concentrations, reactants, products, rate constants, and lambda functions. Access to dictionary contents allows for automatic declaration of symbolic parameters for symbolic computation and easy establishment of inputs for numeric computations. Most numeric computations are handled by NumPy/SciPy [34,35] and symbolic computations by SymPy [36]. We have developed the necessary codes that prepare the inputs for solving ordinary differential equations for integration with SymPy and NumPy modules. We have also implemented functionalities such as tau-adaptive Euler, tau-adaptive Runge-Kutta (RK), fixed interval RK, stochastic simulation algorithm (SSA), tau-leaping algorithm, tau-adaptive chemical Langevin algorithm, fixed interval chemical Langevin algorithm, numerical and symbolic linear noise approximation, and Monte-Carlo expectation maximization, etc. that NumPy/SciPy and SymPy do not have built-in functions.

A relatively new method, network localization [37], allows for studying the qualitative effect of parameter perturbation from structural topology alone. This can be used to obtain physical insights even without propagating the ODE of the system. In BioSANS, both symbolic and numeric network localization is supported.

A list of the codes in BioSANS with a description of its role is provided in sections 7.1 to 7.2 of the supplementary material. BioSANS installer and the actual codes can be downloaded from the following GitHub repositories; https://github.com/efajiculay/BioSANS_installers, https://github.com/efajiculay/SysBioSoft/tree/main/BioSANS

### 3.1 Model construction

To overcome the potential barriers in constructing a model, we have designed a new model input form, the topology file, which is a text file that follows how elementary chemical equations [38] are written. With an example shown in Fig 2, such a file is intuitive to construct and contains only a list of reactions, rate constants, initial conditions, and additional parameter settings. The users do not necessarily need to provide expressions for fluxes or propensity, which is definitely needed when writing a regular code for simulation. BioSANS can automatically infer the propensity from the chemical reaction listed. Algebraic expressions for concentration and propensity modification are supported in case the needed expression differs from mass action kinetics, such as the commonly used Mechaelis-Mention kinetics or the Hill function. Time-dependent propensity and complicated conditional logic (i.e., events, events with delay, etc.) is supported in the algebraic expression. Models can also be written as differential equations in the topology file.

**Fig 2.**
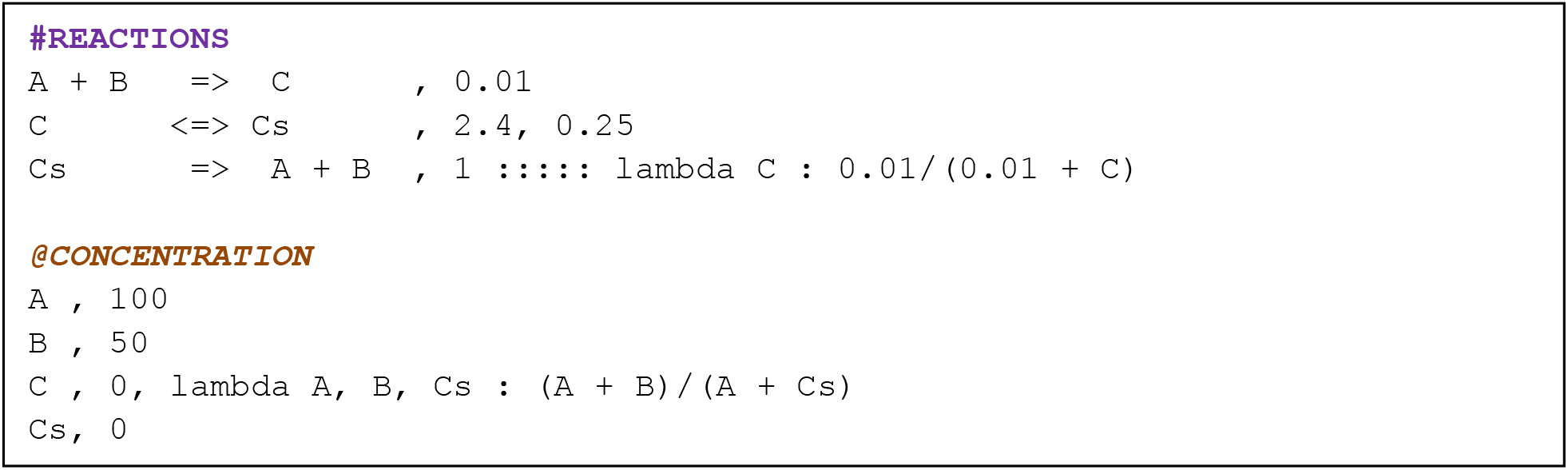
An example of a topology file that can be used to perform simulations in BioSANS.

The BioSANS topology file has three main tags: Function_Definitions:, #REACTIONS, and @CONCENTRATION. Fig 2 shows the last 2 tags, which is the minimal requirement. The single-headed arrow represented as “=>” is used for forward reaction and the double headed arrow “<=>” is for reversible reaction. This setting allows a user to easily identify reactions, one way or reversible, in a topology file. The numbers after the comma in each row after the reactions are the rate constants. The number of rate constants listed should match the arrow used: 1 entry for one-way and 2 entries for reversible reactions. The initial concentration is defined after the comma in each corresponding species under the @CONCENTRATION tag. Propensity and concentration modifications can be written as a lambda expression in each row after some delimiter. If algebraic modifications are provided, the rate constant and or initial concentration will be ignored and the propensity expression is evaluated. The full details of how to construct a topology file including algebraic modifications is in section 4 of the supplementary material. Topology files intended for parameter estimation is discussed is sections 8.5 to 8.7.

Most available models in the literature are provided as a Matlab or Python script or are available in SBML format. BioSANS can run a Python file directly, and ODE models written in Python will be easy to simulate in BioSANS. Shown in Fig 3 is an alternative input file called an ODE file. It can also be used to create models that require only the ODE expression, a set of initial conditions, and a rate constant.

**Fig 3.**
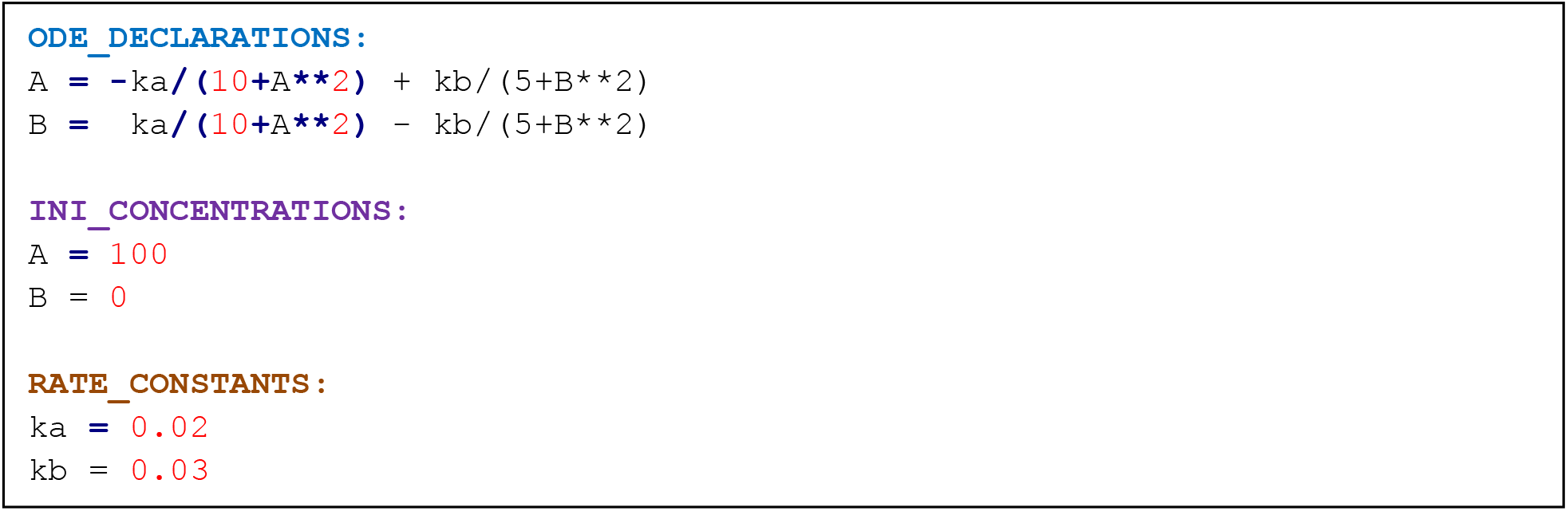
An example of an ODE file with 2 chemical species.

An ODE file can be constructed with minor modification from an existing ODE model and is suitable to use if we have an idea of the underlying mathematics that the system obeys. The ODE_DECLARATIONS: tag is simply the set of species in the left-hand side and the corresponding differential expression in the right-hand side. This file needs to be converted to a topology file with BioSANS before we can propagate the model in the usual way. The converted file will contain the corresponding chemical reaction that satisfies the ODE expressions. Unlike the topology file, the ODE file does not support a lambda expression, and if a user wants to introduce events, such events have to be incorporated in the converted file. SBML files to topology file conversion is also supported in BioSANS for models that are available only in SBML format.

BioSANS also takes line commands with highly human readable formats, similar to SQL commands. Fig 4 shows a model using such commands, which we call SSL. This function allows for automations by saving many SSL scripts in one file and loading them via the BioSANS console. The model in the script is also converted by BioSANS to topology format and executes the remaining statements, which tell BioSANS how to process the model from propagation to analysis.

**Fig 4.**
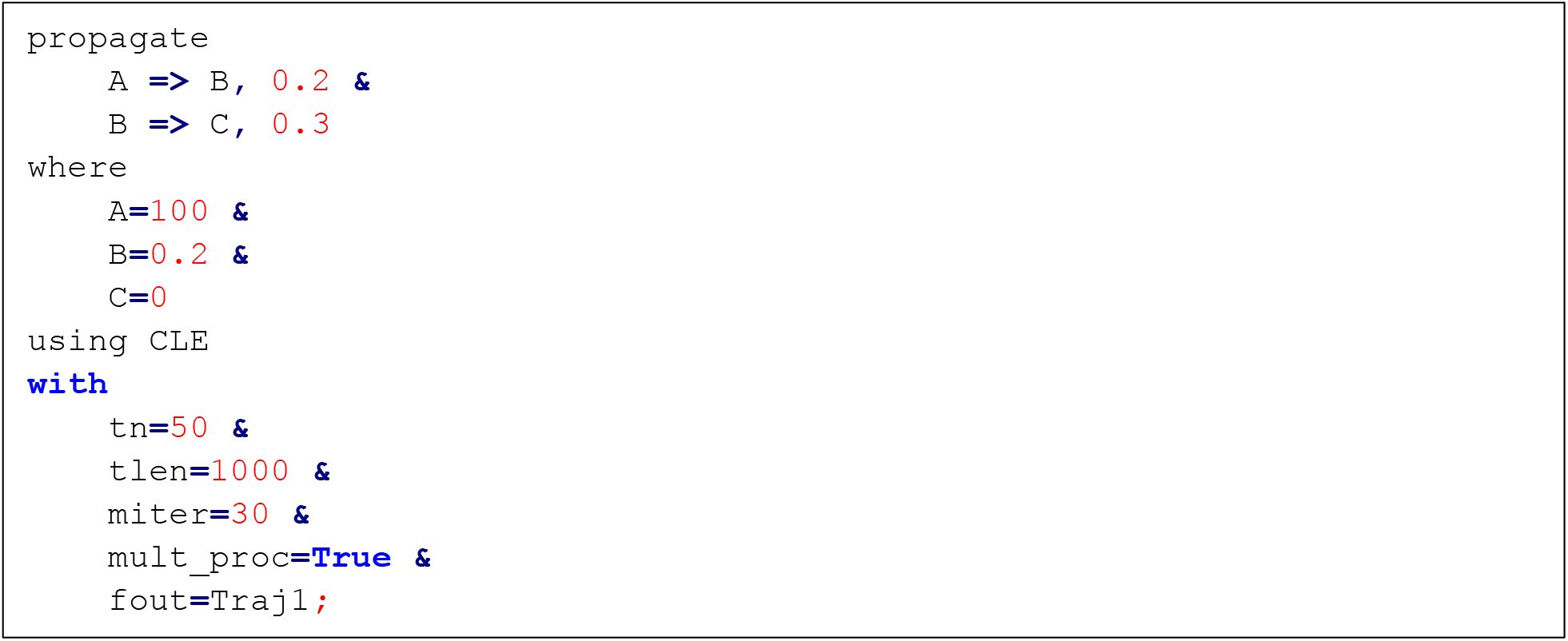
An example of BioSANS structured simulation language (SSL) scripts.

As part of the general model construction, BioSANS can also perform parameter estimation for a given topology file and experimental data, a feature not available in many similar software packages. A tutorial is included in supplementary material for a detailed description of the different BioSANS input files (section 4). Topology files for parameter estimation with examples are discussed in sections 8.5 to 8.7.

### 3.2 Propagation

The trajectory of the state variables can be computed via deterministic and stochastic settings. For deterministic computation, symbolic analytical expression and numerical integration are supported. The stochastic calculation may be carried out with the SSA [39], chemical Langevin equation [40], tau-leaping algorithm [41], and LNA [30,32].

Symbolic computation is currently limited to mostly linear differential equations and a few nonlinear differential equations. The symbolic test performed in this work involves only up to 10 interacting chemical species; beyond that, the software may take a very long time for an answer. Topology files with a modified functional form of propensity with nonlinear functions are likely to fail in symbolic computation. Such capabilities can be further improved if the symbolic ODE solving is improved with SymPy in the future. Nevertheless, our work allows for a quick input in topology and a symbolic answer for solvable systems without the need to declare variables and write ODE expressions. To the best of our knowledge, this is a novel functionality among current systems-biology packages and a feature that is very convenient to use.

In propagating deterministic trajectories, we took advantage of the LSODA [42] algorithm in the SciPy module of python, which allows for fast and efficient numerical integration. We also provide 2 different Euler propagations [43], both with an adaptive time-step setting. Runge-Kutta fourth (RK4) is also offered in 2 version, one is time adaptive and and the other is fixed time-interval [44].

Euler and RK4 were manually coded to support certain SBML features that do not fit the standard use of the LSODA library. The critical SBML features that require coding new integrators are events with delay, events triggered by time, events triggered by other events, use of infinity symbols, and use of SBML keywords such as csymbol delay, csymbol time, rateof, etc. For sophisticated events, it is necessary to keep track of the previous trajectories, so the standard built-in integrator in Python was inapplicable.

Euler propagation is simple and can provide a well-grounded side-by-side comparison with the Euler-Maruyama propagation for stochastic simulation. In our implementation, 2 different adaptive time-step Euler schemes are offered. The first scheme, labelled “Euler (tau-adaptive-1),” was inspired by the tau-leaping scheme [41], with the step size determined by limiting the largest change among the reactions. The second scheme is labelled “Euler (tau-adaptive-2),” which involved choosing a step size that maintains the error as compared with a second-order RK estimate.

RK4 is one of the most popular and widely used integrator. It is simple, accurate in most cases, and easy to implement. Further details about our Euler and RK4 implementation are provided in supplementary material, section 10.2.1.

Stochastic trajectories can be propagated by using our implementation of SSA, 2 different implementations of tau-leaping, and 2 different versions of chemical Langevin algorithms. The SSA, or the Gillespie’s algorithm [39], is an exact realization of the chemical master equation, with probabilistic reactions taking place at the microscopic level. Tau-leaping and chemical Langevin algorithms are inexact stochastic algorithms that accelerate propagation by using a large time step. They allow for many incidents of chemical reactions by using a random variable to account for the number fluctuation.

Our first implementation of tau-leaping, labelled “tau-leaping-1,” includes a regular Poisson random variable describing the number of times a reaction channel fires. In this implementation, the step size was determined by requiring a small change in the reaction propensity following the new tau-selection procedure [41]. Treatment of critical reactions, which are reactions that are close to exhausting some of its reactants after several fires, is not considered in “tau-leaping-1”. We simply draw another random variable when the species concentration becomes negative because this is a very rare event under the new tau-selection procedure. The second version, labelled “tau-leaping-2,” adopts the “modified Poisson tau-leaping,” which separates the treatment of critical and non-critical reactions and also uses the new tau-selection procedure [41]. Critical reactions are monitored and treated as discussed in section C of [41].

The two different versions of Chemical Langevin equations (CLEs) [40] we implemented in BioSANS are the tau-adapted version, labelled “CLE-tau-adaptive,” and a regular fixed-interval version, labelled “CLE-fixed-intvl”. CLE-tau-adaptive employs a simple tau selection we developed, which is fast for non-stiff to moderately stiff problems. (A detailed description of this algorithm is in section 10.2.4 of supplementary material). CLE-fixed-intvl is a typical Euler-Maruyama [45] propagation with a fixed time step.

It is important to estimate the variation in stochastic simulations. In addition to simply analyzing variances and covariances via stochastic trajectories, LNA [30,32] is a convenient way to estimate these quantities of a system. For LNA, we provide both symbolic and numeric ways of solving the covariance matrix. Steady-state LNA and time-dependent LNA computation are supported. Propagation of covariance and Fano-factor is possible in the time-dependent LNA.

BioSANS can directly run python scripts. The script may contain symbolic and numeric models taking advantage of SymPy and NumPy libraries. All functions in BioSANS itself can be used inside a code for customized propagation of trajectories as needed.

### 3.3 Analysis

The analysis part of modeling involves analysis based on topology and post-processing of trajectories. The results of analysis can be used for interpretation and comparison with a known ground truth. BioSANS supports covariance, Fano-Factor, cross-correlation, overall trajectory density plot, density plot binned with time, histogram slice at selected time range, average of trajectory plot, phase portrait, and custom plots. Network localization [37] is also available as an analysis method and can be directly applied to a topology file. Export of a trajectory to a file and plotting is by default enabled after every simulation but can be disabled when needed. Customized analysis is also possible because BioSANS can run Python files.

### 3.4 Testing BioSANS algorithms

The BioSANS algorithms presented above were tested for systems of various complexities and sizes. Performance was qualitatively assessed based on ease of use and features supported. Here we also report 4 quantitative tests of various features of BioSANS: 1) the semantic test for correctly interpreting SBML, 2) the stochastic SBML test suite for the stochastic simulations, 3) tests for symbolic solutions, and 4) tests for parameter estimation.

Scripts for automated evaluation against all tests are used except for symbolic LNA, for which each test is performed manually. The details of the tests are discussed in the following subsection.

#### 3.4.1 SBML sematic test

For the SBML sematic test, we used the fourth-order RK (RK4; implicit output) algorithm, and the test cases are as in [46], consisting of 1780 cases. Those test cases measure the ability of a software to interpret SBML files. We adopt the criteria provided with the sematic test as follows:

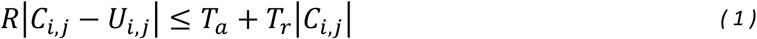

where ***C***_***i***,***j***_ and ***U***_***i***,***j***_ are the expected and estimated value for observable ***i***, at time index ***j, T***_***a***_ is the absolute tolerance for a test case, and ***T***_***r***_ is the relative tolerance for a test case. The details of these tolerance values are included in each test case settings file together with the semantic test cases.

In the semantic test, we added a tau-scaler (step size modifier) in RK4 and reduce this scale sequentially from 0.1, 0.01, 0.001, and 0.0001. If it passes the test, the loop is halted and reported as passed.

#### 3.4.2 The stochastic SBML test

For the stochastic SBML test, stochastic algorithms such as SSA, tau-leaping, and CLE are tested by using the SBML discrete stochastic test suite, containing 39 cases [47]. The suite measures the ability to perform stochastic simulation with less emphasis on interpreting SBML. We adopt the criteria for the mean and standard deviation provided with the test cases.

In the mean test, a scaled z-score is represented as follows:

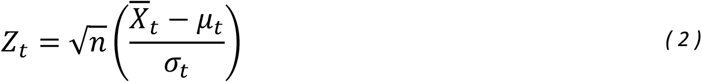

where ***Z***_***t***_ is the z-score at time ***t***, 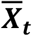 is the mean of trajectories at ***t, μ***_***t***_ is the true or accepted mean at ***t, σ***_***t***_ is the true or accepted standard deviation at ***t***, and ***n*** is the number of trajectories. If ***Z***_***t***_ is in the range (−3,3), the test passes at time ***t***; otherwise it fails.

The standard deviation test makes use of a scaled ratio of the variance, which can be summarized as follows:

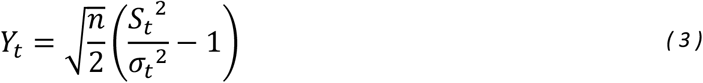

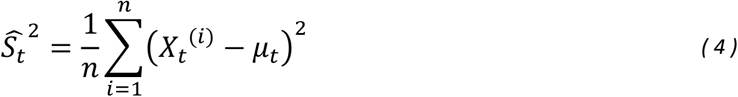

where ***Y***_***t***_ is the scaled ratio of the variance, ***Ŝ***_***t***_^**2**^ is the variance, and ***X***_***t***_^(***i***)^is the value of species ***X*** at time ***t*** for trajectory ***i***. If ***Y***_***t***_ is in the range (−5,5), then the test passes at time ***t***; otherwise it fails.

For the inexact tests, we adopt the mean ratio and standard deviation ratio as suggested in the SBML stochastic test suite for the inexact simulator. We used the 0.95 to 1.05 cutoff for the mean ratio and standard deviation ratio for an acceptable test.

In the stochastic (except CLE) tests, our script reruns the test at most 3 times and it needs to pass at least once to be considered passed. For CLE, we use up to 7 repeats with decreasing tau-scaler. This tau-scaler modifies ***τ*** in both fixed and tau-adaptive CLE. If CLE passes one of the settings, it is considered passed. This is because a randomly chosen tau for CLE may not satisfy the requirement that ***τ*** be small enough for no appreciable change in propensity to occur and large enough that the expected number of occurrences of each reaction channel in the interval is > 1.

#### 3.4.3 Tests for symbolic solution

To test performance on symbolic computation, we created 40 analytical expression test cases and 20 symbolic LNA tests. We used relative absolute deviation (RAD) at each time point from the numerically propagated trajectory as a criterion, defined as follows:

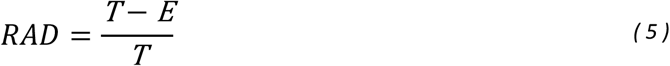

where ***T*** is the true value of the observable, and ***E*** is the estimated value of the observable. ***RAD, T*** and ***E*** can be time-dependent (i.e., if the observable is a species trajectory). If RAD is > 0.05, we consider that it failed the test; otherwise it passed.

For symbolic LNA (covariance) and steady-state concentration, we use numerical results as the true value and RAD to decide a passed or failed test.

#### 3.4.4 Tests for parameter estimation

To test performance on parameter estimation, 45 parameter-estimation test cases were created. The first 40 cases are the same test cases from the symbolic test and the additional 5 cases are for extremely different orders of rate-constant magnitudes. The same RAD as defined in Eq. (5) was used for comparing the rate constants to the true values.

For the parameter tests, failed test cases are manually rerun using different settings for at most 3 times before we finally label the performance on that test case.

## 4.0 Results and Discussion

### 4.1 Ease of use

BioSANS can be used via a GUI, command line interface, and as a Python import, which we believe is a great improvement over other software because many software packages for systems biology require basic programming and commands [21–27]. Some can be used without coding but have limited functionality and computation power if only using the GUI [29].

In handling reactions with their mathematical expressions (as fluxes and propensity), unlike many previous programs, BioSANS takes an intuitive, simple-to-construct topology representation. In most, this topology file, as introduced in section 3.1, does not require declaration of variables and typing propensity expression. Once a topology file is prepared, most of the features available in BioSANS can be used. BioSANS also offers interconversions of topological files and SBML.

The input files for most existing systems biology software requires declaration of the full propensity expression. Some software packages require transforming reversible reactions into 2 forward reactions [21]. Others only accept SBML files as input, which is difficult to construct manually [48]. Software packages for preparing SBML files [49,50] do exist, but newcomers to the field will have to study several before starting to work on the problem they want to simulate.

### 4.2 Features supported as compared to some selected software

In Table 1, we list features supported by BioSANS and some selected software with similar purposes: Copasi [29] Cerena [28], Stochpy [21] and Stochkit2 [27]. In this list, only BioSANS supports symbolic computations. As far as we know, no existing software supports symbolic computation without the need to declare variables and write ODE expressions. Currently, this new feature is available for time trajectories for the species, LNA covariance matrices, steady-state concentration, and network localization.

**Table 1.** Feature comparison between BioSANS and selected software with a similar purpose.

| Feature | Copasi | Stochkit2 | Stochpy | Cerena | BioSANS |
| --- | --- | --- | --- | --- | --- |
| **Numerical analysis** |  |  |  |  |  |
| Exact SSA | * | * | * | * | * |
| Inexact SSA | * | * | * | * | * |
| Linear noise approximation | * |  |  | * | * |
| Parameter estimation | * |  |  | * | * |
| Network localization |  |  |  |  | * |
| **Symbolic analysis (Analytical expression derivation)** |  |  |  |  |  |
| Species analytical expression |  |  |  |  | * |
| Steady state concentration |  |  |  |  | * |
| Linear noise approximation |  |  |  |  | * |
| Network localization |  |  |  |  | * |
| **Simulation options** |  |  |  |  |  |
| SBML support (stochastic) | * | ? | * | * | * |
| SBML support (semantic) | * |  | ? | * | * |
| Easy to prepare input<sup>a</sup> |  |  |  |  | * |
| Actual time steps output |  | ? | * |  | * |
| Fixed-interval output | * | * | * |  | * |
| **Output analysis** |  |  |  |  |  |
| Auto-correlation |  |  | * |  | * |
| Histogram distance |  | * |  |  |  |
| Propensities | * |  | * |  |  |
| Moments |  |  | * |  |  |
| Waiting times |  |  | * |  |  |
| Probability density with time |  |  |  |  | * |
| Time-slice of densities |  |  |  |  | * |
| Average of trajectory |  |  |  |  | * |
| Bootstrapped covariance |  |  |  |  | * |
| **Software characteristics** |  |  |  |  |  |
| Plotting | * | ? | * |  | * |
| Data exportation | * | * | * |  | * |
| GUI | * |  |  |  | * |
| console | * | * | * |  | * |
| Flexible environment | * |  | * |  | * |

a - do not necessarily need to type rate law expression, no need to mapped variables
? - feature not fully supported with limited capability or not supported at all
SSA, stochastic simulation algorithm; SBML, systems biology markup language; GUI, graphical user interface

Table 1 also shows that BioSANS features a GUI, an easy input format; parameter estimation; network localization; and LNA. Some software packages provide limited SBML support (unquantified) and others do not claim support at all. BioSANS supports all features listed except for histogram distance, propensities, moments, and waiting times in the output. Those exceptions can be calculated from trajectory data, and because BioSANS can run Python script, an advance user can also calculate those unsupported features.

As an important implementation, the topological input of BioSANS supports coupled propagation of dependent and independent systems in one file. This can help mimic a mixture of unrelated chemical reactions that coexist. This feature works for both deterministic and stochastic settings except for SSA and tau-leaping2.

COPASI is a popular and powerful software in systems biology that supports many types of computations. It is widely used in various fields and constantly maintained by a group of developers. It is relatively easy to use with a GUI, console interface, and bindings to other scripting languages. As compared with BioSANS, it does not support symbolic or analytical expression. The GUI in COPASI does not support multiple trajectory sampling (ensemble of trajectory sampling) for stochastic simulations. Model creation in the GUI requires typing the rate law if it is not present in the list of functions or not previously saved. It supports models imported from SBML, but how much support it provides to the SBML stochastic test suite is not clear. The console interface and bindings it provides can be used together or called in other programming languages such as Python, c/c++, etc. but is usually critical with versions. Together with some non-trivial programming skills, COPASI is a powerful software for systems biology.

Stochpy is a Python-based software that is dedicated to stochastic simulation. It can make use of Python libraries, which can be very powerful to use. It is the only software that passed the SBML stochastic test suite when it was published. Model creation is easy and follows the PySCeS model description language, but it requires typing the rate law expression. It can import SBML files, but the SBML semantic test results are not seen on the SBML website. It provides support for fixed-interval output and actual-time output. However, as a Python module, it can only be used via command line and as a library import. Moreover, as compared with BioSANS, Stochpy does not provide support for symbolic computation, parameter estimation, LNA, and deterministic calculation.

Stochkit2 is a software for discrete stochastic simulation, its main advantage being speed and parallelization. Models can be created in an xml-like format using certain tags and also allowing for defining arbitrary function. Its latest version released support events (conditional statements) as well. However, Stochkit2 does not quantify how much SBML support it provides. As compared with BioSANS, the rate law expression is not automatically inferred and requires typing under the rate tag. It does not provide support for GUI, parameter estimation, and symbolic computation nor theoretical computation such as LNA.

CERENA is a MATLAB-based software that provides many functions. It supports time-dependent propensity, parameter estimation, LNA, and other high-level mathematical analysis. Models can be created using a model definition file, which uses MATLAB-like declarations. It supports importing SBML files, which allows for producing the model definition file automatically. Nevertheless, CERENA does not provide support for symbolic computation, does not provide a GUI and can only be used by MATLAB programmers. The model definition file it provides is not as intuitive as chemical elementary equations.

BioSANS supports a wide variety of features as mentioned above and in the previous section. The GUI and console interface (structure simulation language) allow for multiple trajectory simulation and support parallel runs by taking advantage of a multiprocessing library [51] in Python. It supports time-dependent propensities and complex logic in the topology but requires some experience to properly set up complicated logic if necessary. Because it is Python-based, it can also make use of powerful libraries from Python, which will be very useful in automation (i.e., model selection if used as a Python library). Hence, BioSANS is a software that makes systems biology modeling available even to nonexperts and is well suited for newcomers in the field. It can be used for teaching systems biology and for performing simple to complex modeling tasks.

### 4.3 Performance and testing of the software

The following sub-section summarizes the performance of BioSANS against various tests. The detailed results are provided in section 11 of the supplementary material.

#### 4.3.1 Semantic test

In Table 2, we list the results of SBML sematic tests comparing BioSANS and other software that submit results to the SBML database website. BioSANS passed most of the test cases, with about 70% correct test results. This is already above the average level for SBML support among all the software packages that submit their results.

**Table 2.** Comparison of performance of several software packages for the SBML semantic test case.

| Software<sup>a</sup> | Date submitted | Passed |
| --- | --- | --- |
| BioUML 2018.2 | 18-Jun-18 | 100% |
| Morpheus 2.1 | 18-Jan-19 | 81% |
| **Biostoch/BioSANS1.0** | 7-Dec-20 | 71% |
| BioUML 0.9.5 | 4-Jun-13 | 67% |
| COPASI Build 4.23.189 | 12-Jun-18 | 67% |
| iBioSim 2.4.2 | 4-Feb-13 | 67% |
| Simulation Core Library 1.2 | 28-Mar-13 | 63% |
| libRoadRunner 1.3 | 29-Dec-14 | 60% |
| RoadRunner 2.10.0 | 16-Jan-13 | 60% |
| WinBEST-KIT 2.0.0 | 20-Dec-18 | 60% |
| LibSBMLSim 1.1.0 | 31-Jan-13 | 55% |
| AMICI 0.11.8 | 1-Sep-19 | 20% |
| FluxBalance 1.9 | 21-Jul-14 | 1% |

<sup>a</sup> The software included here are those that submit their results to an SBML website.

Currently only BioUML (Kolpakov *et al*., 2019) provides 100% support for all semantic standards set by SBML. BioUML is a general package of programs for systems biology that provides a lot of functionality (300+ different types of analysis if connected to Galaxy), including deterministic and stochastic simulation. Its major advantage is being a web-based software, so it is accessible to users who have a BioStore account. It supports analysis of biomedical data from omics experiments, parameter estimation, databased links, etc. It can generate codes in Java from models created via its diagram editor in a drag-and-drop manner. Conversely, BioUML does not provide support for symbolic computation, LNA, network localization and does not mention performance on stochastic SBML tests.

Morpheus is a modeling software that focuses on the study of multi-scale and multicellular systems using ordinary and partial differential equations [52]. Models can be created by describing them in the GUI instead of in a code and the inputs can be symbolic expression. It can plot cellular processes and dynamics in an image, depicting actual cells, tissues, etc. involved in the process. However, even with the GUI, a user still needs to define the global variables, the system of differential equation and other constraints. The creators did not claim to give output in a form of symbolic/analytical expression. The support for parameter estimation is limited to parameter sweeping.

Currently, the SBML support BioSANS offers has some limitations. Features such as overlapping delays, CSymbolDelay, use of infinity symbols, and latest features available only in level 3.1 and above are not fully supported. To interpret SBML files, BioSANS converts the SBML file to a topology file. If BioSANS fails to convert the file properly, there will be an error at run time and no output will be returned. Manual inspection and basic editing of the converted file may help correct the conversion error. Still, this will require understanding the system to properly correct the file or reading the SBML manually to check for inconsistencies.

#### 4.3.2 Performance on stochastic test

BioSANS exact stochastic algorithms are tested by using the SBML discrete stochastic model test suite (SBML DSMTS). In the DSMTS test, a good number (10,000 in our case) of stochastic trajectories was collected for each of the 39 test models, with the standard deviation for the mean and the variance calculated at 50 time points. Table 3 lists the algorithms and the number of test cases with their corresponding percentage of time points passed in the DSMTS tests as columns. Following the suggested routine, owing to the probabilistic nature of such simulations, 2 to 3 error cases in the mean test and 4 to 6 errors in the variance test are expected for a perfectly working stochastic algorithm under the exact SSA test [47]. The performance of our SSA implementation falls within the criteria, with few cases suffering considerable discrepancy mostly in the timed trigger cases, possibly because of the treatment of exact timing and the subsequent error from such an uncertainty. The time-triggered cases are still within the 90% to 99% window if tested under the inexact test. To the best of our knowledge, we cannot find software comparisons reported for stochastic test cases. We have also performed tests with StochPy, which is claimed to have passed SBML DSMTS tests. StochPy and BioSANS show comparable performance for SSA as listed in Table 3. Thus, BioSANS SSA implementation is reliable.

**Table 3.**
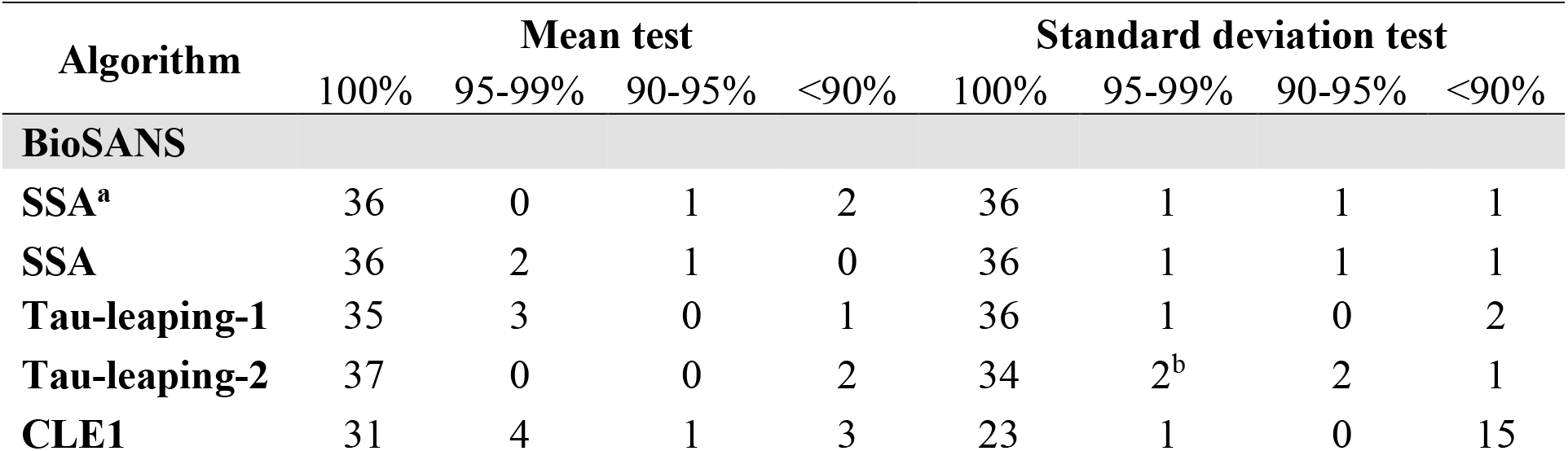
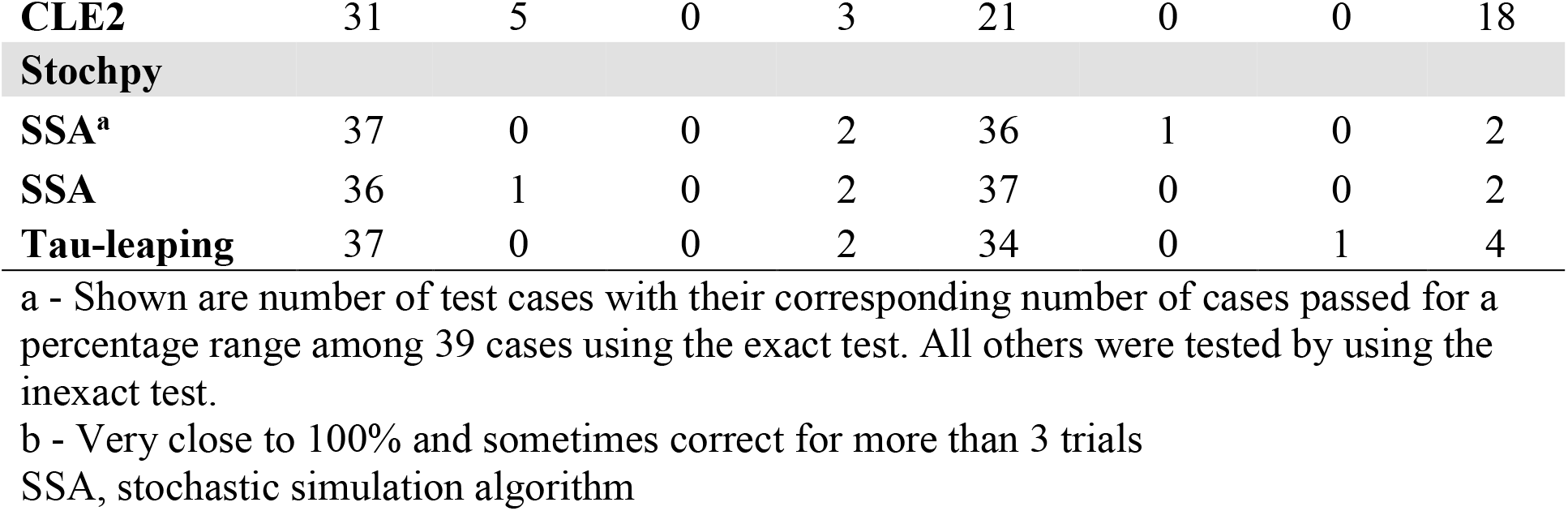
Comparison between StochPy and BioSANS in passing tests in the SBML discrete stochastic model test suite (DSMTS)

The approximate algorithms in BioSANS were also tested by using SBML DSMTS. For approximate simulators, SBML DSMTS suggested observing the ratio of the mean and variance obtained from reasonable sampling trajectories (10,000 in our case) with their corresponding standard values (provided in the test). The inexact tests in Table 3 show the number of cases for a certain percentage with estimated mean or variance within 0.95 to 1.05 of the standard value. Tau-leaping2 has 2 errors in the mean test and 5 errors in the standard deviation test. This is still within the allowed range of error suggested in SBML DSMTS. The data in Table 3 are the best outcome for 3 repeats of 10,000 trajectory simulations. The detailed results of the tests and explanation for the failed results are included in supporting information. A failed result does not necessarily mean it is too far from analytically derived trajectories, and we believe that it is still useful for qualitative purposes. From the inexact test, the BioSANS tau-leaping-2 is comparable to StochPy tau-leaping implementation. CLE1, CLE2, and tau-leaping-1 have slightly more errors than the allowable number, but they are generally faster algorithms. Therefore, these methods are suitable for systems that require a large amount of computation, but they should be used cautiously.

#### 4.3.3 Performance in symbolic tests

The following table summarizes the performance of BioSANS in symbolic computation. Typically, all species in symbolic solution should be a function of time ***t***. It would be desirable for general solutions, with symbolic dependence on the initial concentration ***x***_**0**_, and parameters such as rate constant ***k***. In BioSANS, we provide a way to report analytical expressions in 4 different categories of functions such as ***f(t), f(t***,***x***_***0***_***), f(t***,***k)***, and ***f(t***,***k***,***x***_***0***_). This notation corresponds to functions of the remaining variables (after substitution) before symbolic computation is attempted. We refer to ***f(t***,***k***,***x***_***0***_) as a pure symbolic expression and others are called semi-analytical expressions. The semi-analytical expressions are easier to solve by using SymPy, which has the majority of test cases passed, as seen in Table 4. The pure symbolic expression is not easy to handle and often generates large analytical expressions. Sometimes it also takes a lot of time to finish computation.

**Table 4.**
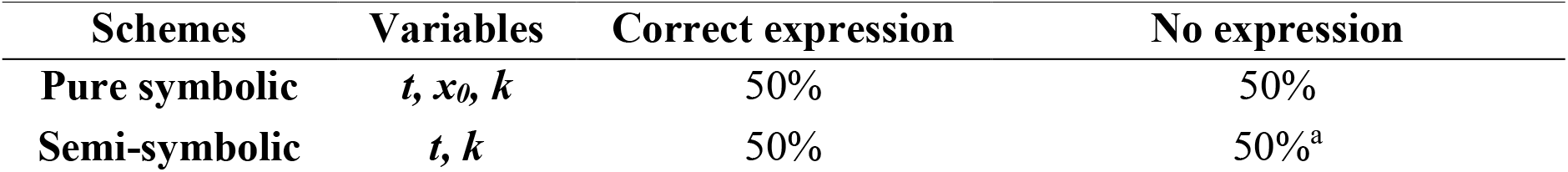
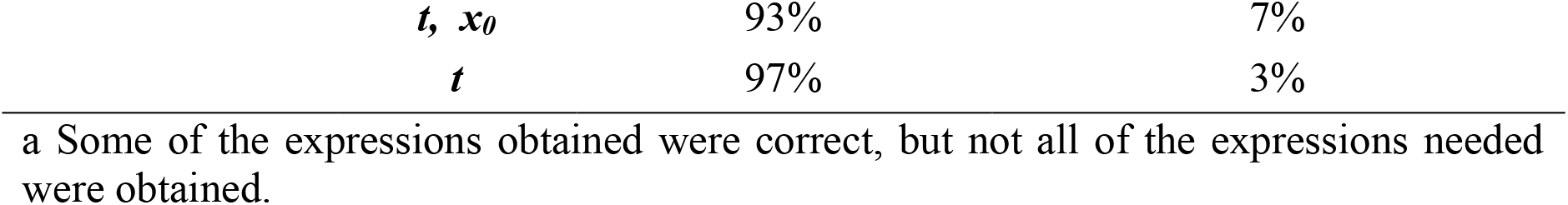
Performance of BioSANS in generating analytical expression for species concentration with different symbolic computation modes.

For the analytical expression, BioSANS supports only linear differential equations and few nonlinear differential equations. There were some linear ODEs that BioSANS could not fully support, such as overlapping reversible reaction, overlapping cyclic and reversible reaction, cyclic structure inside cyclic structure, and overlapping cyclic structure. Some of these not-fully-supported linear ODEs can still work using the ***f(t)*** mode. Our test cases consist of reactions with fewer than 10 chemical species. It is possible to solve systems with greater than 10 species especially using ***f(t)*** and ***f(t***,***x***_***0***_), mode. Other modes will take longer, and the expression might be larger, so not usable for physical interpretation. The ***f(t***,***x***_***0***_) mode describes the effect of initial concentration on the species analytical expression and is easier to solve than the pure symbolic mode. Dependence on the rate constant can be inspected by using the ***f(t***,***k)*** mode, but it is almost as difficult to calculate as with a pure symbolic solution. The ***f(t)*** mode is useful if we only want to know the trajectory with respect to time and do not intend to discover the relevance of the initial concentration and rate constant in the final trajectory.

Symbolic expression with the steady-state LNA for the covariance matrices can be reported in the same 4 modes as well. The steady-state concentration expression is included when computing symbolic LNA. Table 5 summarizes the performance in symbolic computation for the covariance matrix analytical expression and steady-state analytical expression. The None entry in the variables column pertains to a numeric answer but is symbolically derived.

**Table 5.** Performance of BioSANS in generating LNA and a steady-state analytical expression.

| Variables | Correct expression |  | Takes too long |
| --- | --- | --- | --- |
|  | Steady state | Cov/Var |  |
| LNA-symbolic-$f(x_0, k)$ | 95% | 55% | 45% |
| LNA-symbolic-$f(x_0)$ | 95% | 90% | 10% |
| LNA-symbolic-$f(k)$ | 95% | 55% | 45% |
| None<sup>a</sup> | 95% | 90% | 10%<sup>b</sup> |

a Numeric answer but symbolically derived
b Some returns wrong expression

The symbolic computation functions in BioSANS are currently useful for small- and moderate-sized systems. We think it necessary to have symbolic capability because analytical expression can give more physical insights than can simulations. Whenever a system can be simplified to a size whereby we can generate an analytical expression, our BioSANS symbolic computation can help generate a reasonable expression.

Symbolic solving ODEs with programs has become more available in recent years. As the development of SymPy progresses, BioSANS symbolic solving could be further improved.

#### 4.3.5 Performance in parameter estimation

The test for parameter estimation is assessed (using RAD) by the ability to reproduce the trajectory and rate constant values. Simulated trajectories are generated for 45 models with given rate constants. Most of the simulated trajectories contain 201 data points, except for test cases 15, 19, and 43, with 1000 data points. Test cases 15 and 19 include reversible reactions, and test case 43 is a stiff problem. The algorithms in Table 6 are used to estimate the rate constants given all species concentrations as a function of time. Even though this is an ideal case because most biological experiments allow for measuring only a few numbers of species and with limited time points, our test for BioSANS aimed to provide the minimal requirement for estimating parameters for mathematical modeling.

**Table 6.** Performance of different algorithms in BioSANS for parameter estimation.

| Algorithms | Speed<sup>a</sup> | Rate constant | Trajectory |
| --- | --- | --- | --- |
|  |  | Percent passed |  |
| **Nelder-Mead (NM)** | Fast | 93% | 98% |
| **Monte Carlo EM** | Slow | 93% | 93% |
| **Differential evolution (DE)** | Slow | 93% | 93% |
| **NM-DE-NM combined** | Slow | 98% | 98% |
| **Powell** | Fast | 91% | 91% |
| **L-BFGS-B** | Fast | 80% | 87% |
| **Parameter slider<sup>b</sup>** | N/A | 44% | 64% |

a A rough estimate of the time required. Usually “fast,” within 5 min, with a typical desktop computer. (slow ≥ 30 min)
b A graphical user interface element for sliding values of parameter

Table 6 summarizes the performance of BioSANS parameter-estimation algorithms given all the species trajectory. Most of the algorithms can capture the true rate constant and trajectory except for the Newton-based algorithm, L-BFGS-B. See section 11.5 of the supplementary material for a detailed classification of test results. There is always a chance of being insensitive to some of the parameters in the dynamics of a system, and parameter estimation based on the trajectories is subject to this natural constraint. Therefore, even if some rate constants are not close to the true value, the trajectories can still be reproduced.

Insights into parameter perturbation is easier to interpret visually. BioSANS offers a parameter slider, which is a GUI element that allows for visually inspecting perturbation effects. With this feature, a user can modify parameters and see the effect of the modification in the results shown as plots. Given some plotted experimental data, we can slide the parameter such that the results of the ODE are similar to those of the experimental data. This will allow for some insights into the parameter in a qualitative sense. However, this is only useful for small systems; as we can see in Table 6, it is the least performant.

### 4.4 Other functionalities

The law of localization in chemical reaction networks, known as network localization, mentioned in section 3.0, is a useful way to investigate perturbation effects of a network based on only topological information [37]. BioSANS supports symbolic and numeric computation of network localization in the form of network sensitivity. This is a qualitative alternative to symbolic solution of species concentration and covariance when SymPy fails to give an answer. From network localization, one can infer the effect of perturbation of a rate constant to species steady-state concentration and covariance. However, we did not provide a test case for these functionalities.

## Conclusion

BioSANS is a software for both experts and nonexperts that supports most of the tasks needed in systems biology modeling including useful features left out by other software. The symbolic computation capability in BioSANS provides analytical expression of solvable cases without the need to type the ODE expression and declaring variables. From model creation, propagation, and analysis, BioSANS provides reliable algorithms that can facilitate the modeling process.

BioSANS can be extended to other fields that make use of ordinary differential equations. For example, ODE is heavily used in pharmacokinetics, pharmacodynamics, Epidemiology, environmental models, economic models etc. ODE is also used in studying reaction mechanisms in chemistry. As long as we can represent the state of the systems as symbols that follow ODE, BioSANS may be used to model such systems.

## Acknowledgement

We express our gratitude to Mr. Yu-Chuan Chen and Dr. Yi-chen Chen for using BioSANS and providing suggestions on how to make it better.

## Supporting Information

**S1 File. Supplementary Information for BioSANS:** A Software Package for Symbolic and Numeric Biological Simulation.

## Notes

### Competing Interest Statement

The authors have declared no competing interest.

